# Dual-modality, deep-learning-enabled endomicroscope with large field-of-view and depth-of-field for real-time in vivo imaging of epithelial hallmarks of cancer

**DOI:** 10.64898/2026.01.14.699592

**Authors:** Huayu Hou, Jimin Wu, Jinyun Liu, Vivek Boominathan, Argaja Shende, Karthik Goli, Jennifer Carns, Richard A. Schwarz, Ann M. Gillenwater, Preetha Ramalingam, Mila P. Salcedo, Kathleen M. Schmeler, Tomasz S. Tkaczyk, Jacob T. Robinson, Ashok Veeraraghavan, Rebecca R. Richards-Kortum

**Author notes:** These authors contributed equally (Huayu Hou, Jimin Wu).

## Abstract

In vivo microscopy (IVM) has shown great promise to improve early detection of epithelial precancer, but it suffers from fundamental trade-offs that limit the resolution, field-of-view (FOV) and depth-of-field (DOF). Here, we present PrecisionView, a compact, deep-learning-enabled endomicroscope that breaks these constrains and achieves 20 mm^2^ FOV and 500 µm DOF with 4 µm resolution, representing approximately 5× increase in FOV and 8× larger DOF compared to conventional IVM with similar resolution. PrecisionView integrates a deep-learning optimized phase mask and real-time reconstruction, enabling rapid in vivo assessment of two key hallmarks of cancer: epithelial cell nuclear morphology and subsurface microvasculature through fluorescence and reflectance imaging. By imaging oral cavity of healthy volunteers and cervical specimens with precancerous lesions, PrecisionView generates large-scale (1-3 cm^2^) co-registered maps of cellular and vascular structures, revealing distinct microscopic patterns associated with anatomic structures and precancerous lesions. Our results suggest the potential of this computational endomicroscope to address the unmet need for early cancer detection at the point-of-care.

## Introduction

Cancer is a leading cause of premature death worldwide^1,2^, and epithelial cancers account for 80-90% of all cancer cases^3^. Most epithelial cancer patients are diagnosed with late-stage disease due to a lack of effective diagnostic methods, resulting in low overall five-year survival rates^4,5^. Histopathologic examination of tissue biopsy is the gold standard to diagnose dysplasia and cancer. However, biopsy is an invasive procedure and requires substantial infrastructure to prepare and interpret histology slides. In the case of large, heterogeneous lesions, only limited tissue areas can be assessed with biopsy. Because most suspicious lesions are benign^6,7^, it’s extremely challenging even for experts to determine where and when to perform invasive biopsy^8,9^. As a result, the accuracy of pathologic diagnosis suffers from sampling error^10,11^.

In vivo microscopy (IVM) has shown promise to improve the early detection of epithelial cancers^11–13^. The morphological hallmarks of cancer include alterations in neoplastic cell nuclei and supporting microvasculature provide predictive hallmarks of precancer^9,14–20^. Simultaneous assessment of changes in both cell nuclei and microvasculature can potentially improve accuracy for early cancer and precancer detection. Pilot studies have demonstrated promising clinical performance of IVM for a variety of cancer types^9,18–20^. However, clinical translation of IVM is hindered by fundamental limitations of conventional optics. The field-of-view (FOV) of current IVM is too small (usually < 0.5 mm^2^) relative to typical size (∼cm^2^) of suspicious lesions^19–32^. The scale-dependent geometric aberrations from miniature optics^33^ or the small size of image relay probes^20,21,23–25,30^ fundamentally restrict the FOV of imaging systems. It is difficult to fully examine large, heterogeneous lesions at risk with current IVM to identify which regions may harbor focal neoplasia^11^. Expanding the FOV with conventional approaches dramatically increases the system form factor and complexity, reducing clinical usability. Integrating lateral scanning and image mosaicking provides an alternative approach to overcoming FOV limitations. However, the limited capture area per image acquisition makes it challenging to maintain sufficient overlap during in vivo tissue imaging. As a result, the total mosaicked imaging field is usually restricted to less than 10 mm^2^, with missing regions caused by gaps in the imaging path^21,34,35^.

Another challenge is that the depth-of-field (DOF) of current systems is inherently coupled to the numerical aperture (NA) and therefore spatial resolution. To achieve cellular resolution for evaluation of microscopic features (< 4 µm), conventional microscopes typically have a DOF less than 60 µm^33,36,37^. This limited DOF is insufficient to accommodate irregularities in the tissue surface, which can be up 200 µm^36–38^, resulting in partially out-of-focus images and loss of critical microscopic information. This issue is more significant when imaging across a large tissue area with a large-FOV device. Moreover, current IVM with limited DOF is not capable of simultaneously capturing in-focus images of both cell nuclei and microvasculature. While cell nuclei are best visualized at the epithelial surface, supporting microvasculature is located 100-200 µm deeper^20,21^. The depth of microvasculature also varies over a large depth range of several hundred of microns^20^. Addressing these differences in axial location with conventional systems requires an additional axial scanning module to enable continuous adjustment of the focal plane, which significantly increases cost and complexity, and reduces clinical usability.

Computational imaging technologies and deep learning have emerged as powerful tools to overcome the limitations of conventional optics. The integration of wavefront encoding with computational algorithms has proven to be a promising approach for improving system capabilities beyond those of conventional microscopy, enabling extended DOF^36,37,39–41^, aberration correction^42–44^, single-shot 3D imaging^45,46^, and lensless imaging^47–50^. Moreover, recent advances in deep learning and optical fabrication enable a novel end-to-end system design strategy, where optics and algorithms are jointly optimized by a deep neural network^36,37,46,51^. This artificial intelligence (AI)-driven optimization at the system development stage unlocks new possibilities to enhance imaging performance. However, only a limited number of computational imaging systems have successfully demonstrated the ability to resolve cellular structures in dense tissue, and even fewer have achieved in vivo imaging capability.

In this study, we present PrecisionView, a compact, AI-powered handheld endomicroscope designed to address the fundamental limitations of conventional IVM and improve in vivo early cancer detection at the point-of-care. By leveraging a deep learning end-to-end optimization framework, we simultaneously optimized both the phase mask design and reconstruction algorithm using a deep neural network. This framework is tailored to enable both fluorescence imaging of cell nuclei and reflectance imaging of microvasculature with substantially increased FOV and DOF (Fig. 1, Supplementary Fig. 1). The wavefront-encoding phase mask is positioned at the Fourier plane of the system to provide phase modulation, generating spatially- and depth-invariant point-spread-functions (PSFs). The resulting image degradation is then corrected by the reconstruction algorithm, which performs image deconvolution and deblurring to restore fine image details. By integrating deep learning end-to-end optimization and simple off-the-shelf optics, PrecisionView achieves a 5.2 mm × 3.9 mm FOV and a 500 µm DOF with 4 µm resolution, while maintaining a compact form factor and a $3,000 cost of goods. This represents approximately a 5-fold increase in FOV and 8-fold improvement in DOF compared to conventional IVM systems of similar resolution (Fig. 1e). With the 40-fold larger volumetric FOV and the capability of fluorescence and reflectance imaging, PrecisionView enables large-scale, high-resolution, co-registered mapping of cell nuclei and microvasculature without requiring refocusing (Fig. 1c,d). The deep learning reconstruction algorithm provides real-time image reconstruction, which allows immediate microscopic evaluation of suspicious lesions at 15 frames per second (FPS), providing diagnostic guidance at the point-of-care (Supplementary Video 1). We experimentally validated performance of PrecisionView by imaging standard test targets, ex vivo porcine tongue specimens, and post-mortem human breast specimens, demonstrating superior resolution across an extended FOV and DOF compared to a conventional system. Furthermore, we established the in vivo imaging capability of PrecisionView by imaging the oral cavity of healthy volunteers. Through handheld scanning and image stitching, we successfully mapped cellular and vascular features over a tissue area > 1.3 cm^2^ at high resolution, revealing distinct features of human lip mucosa and tongue. To assess PrecisionView’s clinical applicability, we performed ex vivo imaging of freshly resected cervical specimens with precancerous lesions, generating maps covering tissue areas > 2.8 cm^2^ via handheld scanning. The system clearly delineated microscopic features corresponding to anatomical structures and pathological abnormalities. These results highlight PrecisionView as a promising noninvasive, rapid diagnostic tool with potential for identification of early epithelial neoplasia at the point-of-care. Our research establishes a novel design framework for the development of next-generation clinical imaging systems with enhanced performance, addressing the significant unmet need to design imaging systems to improve early cancer detection especially in settings with limited medical resources.

**Fig. 1.**
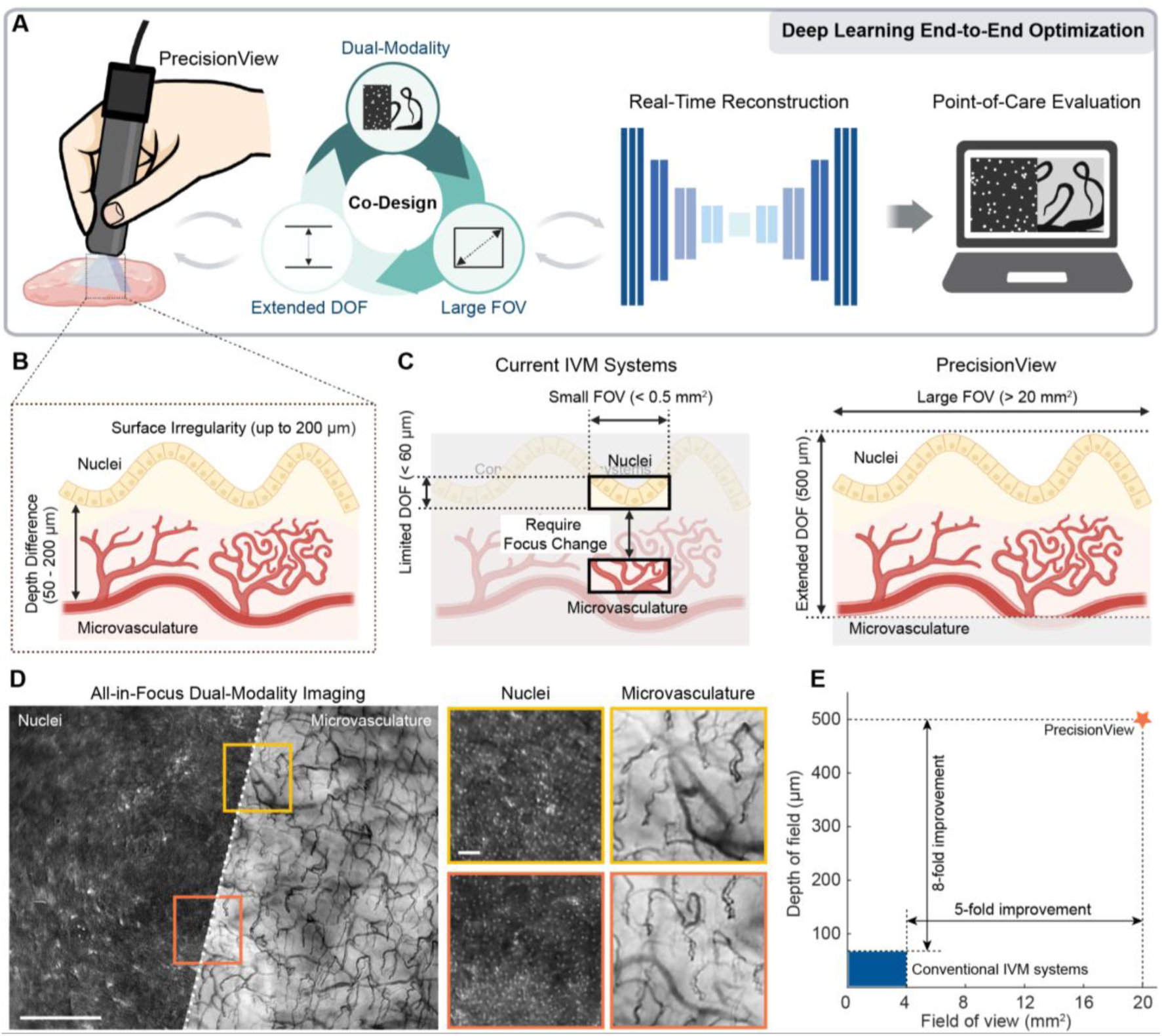
PrecisionView is a compact AI-powered handheld endomicroscope for real-time in vivo neoplasia detection at the point-of-care. **a** Schematic of the design principle and the imaging setup. PrecisionView is a compact, handheld, and simple-to-use endomicroscope designed for high-resolution, noninvasive imaging of epithelial tissue. By leveraging a deep learning framework to co-optimize hardware and software, PrecisionView enables fluorescence imaging of cell nuclei and reflectance imaging of microvasculature, while achieving significantly increased FOV and DOF compared to conventional systems. The deep learning reconstruction algorithm provides real-time video-rate image reconstruction for immediate microscopic evaluation of suspicious lesions, aiding in precancer and cancer detection at the point-of-care. **b** In vivo imaging of epithelial cell nuclei and microvasculature requires accommodation of surface irregularities (axial variations of up to 200 µm) and a broad range of depth differences between epithelial nuclei and subsurface microvasculature (typically 50–200 µm). **c** For existing conventional IVM systems, the FOV is fundamentally restricted which makes it challenging to assess large, heterogeneous lesions at risk. Due to their limited DOF, conventional high-resolution IVM systems require adjustment of the focal position to visualize cell nuclei and microvasculature at different depths. By integrating deep learning end-to-end optimization and simple off-the-shelf optics, PrecisionView overcomes these limitations and achieves an extended FOV and DOF. This enables rapid, co-registered mapping of cell nuclei and microvasculature in a large tissue area at cellular resolution. **d** Representative PrecisionView images of the oral mucosa from a healthy volunteer. The system achieves a 5.2 mm × 3.9 mm FOV and a 500 µm DOF with 4 µm resolution, enabling all-in-focus microscopic visualization of cell nuclei and microvasculature. Scale bars: full image, 1mm; zoom-in views, 100 μm. **e** Comparison of FOV and DOF between PrecisionView and current IVM systems. By incorporating deep learning end-to-end optimization, PrecisionView achieves a 5-fold increase in FOV and an 8-fold improvement in DOF compared to current IVM systems with similar resolution. Without the phase mask and image reconstruction, a conventional endomicroscope using the same conventional optics, demonstrates only a slightly improved FOV with no enhancement in DOF.

## Results

### PrecisionView system design and end-to-end optimization architecture

PrecisionView enables direct, in-focus imaging of different morphological features over a large FOV (5.2 mm × 3.9 mm) and extended DOF (500 µm), utilizing a compact optical layout with end-to-end co-optimization of a phase mask and reconstruction algorithm. The system employs off-the-shelf achromatic lenses and a conjugated optics design, with identical optical designs used for both the microscope objective and tube lens^52^. These components are aligned and housed within a custom 3D-printed enclosure (Fig. 2a). Each of the objective and tube lens assemblies consists of three 12 mm diameter achromatic lenses (Supplementary Fig. 2). A 520/40 nm bandpass emission filter and a jointly optimized phase mask are placed between the objective and tube lens to form the imaging path.

**Fig. 2.**
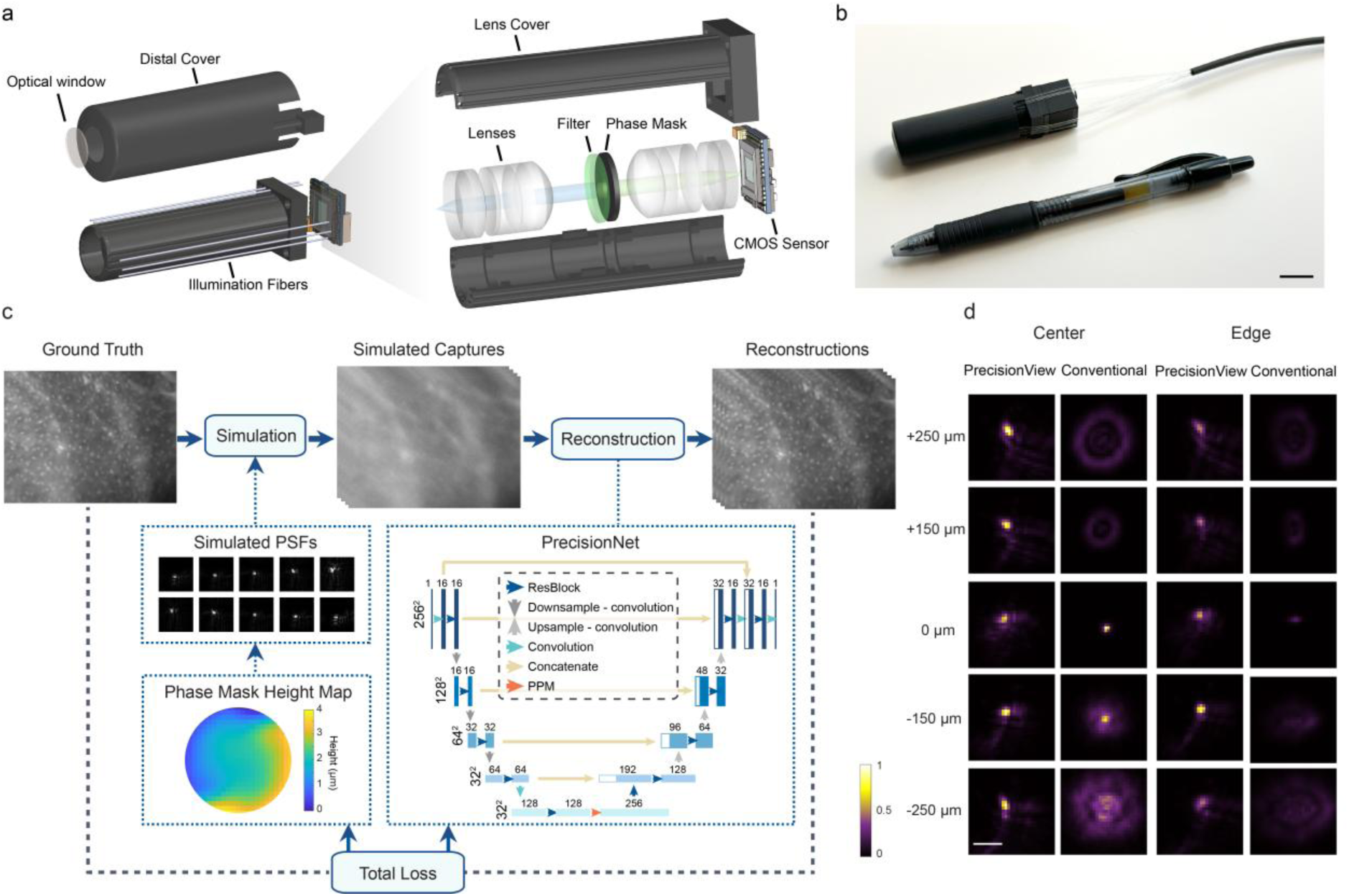
Deep learning end-to-end design of PrecisionView. **a** CAD rendering of PrecisionView. The device incorporates a simple conjugated microscope design with off-the-shelf lenses, a bandpass filter, an end-to-end optimized phase mask, and a miniature complementary metal-oxide semiconductor (CMOS) sensor. The phase mask is positioned at the Fourier plane of the system to enable phase modulation. Fourteen illumination fibers, attached externally to the lens cover, deliver light from a dual-color LED light source. A protective system cover with a sapphire optical window ensures safe contact imaging while maintaining the optimal working distance. **b** Photo of the assembled PrecisionView prototype. Scale bar, 10 mm. **c** End-to-end deep learning framework for joint optimization of the phase mask and reconstruction algorithm. At the optical layer, the end-to-end network simulates the PSFs encoded by the phase mask, generating simulated image captures within the targeted DOF. These simulated captures are then processed by the sequential reconstruction network (PrecisionNet), producing in-focus images. The end-to-end network simultaneously optimizes both the phase mask design and the reconstruction algorithm by minimizing the loss between the network output and ground truth. **d** Experimentally captured PSFs of PrecisionView compared to a conventional system without the phase mask, across the 500 μm depth range at the center and edge (2.5 mm from the center) of the FOV. The trained phase mask enables PrecisionView to generate spatially- and depth-invariant PSFs, facilitating the network digital layer to perform image reconstruction and extend the FOV and DOF. Scale bar, 20 μm.

System illumination is provided by 14 optical fibers symmetrically positioned around the lens housing. The fibers are bundled together and connected to a light source containing blue and green light-emitting diodes (LEDs), The two LEDs are paired with appropriate filters and combined via a dichroic mirror to enable rapid switching between blue and green illumination for fluorescence and reflectance imaging, respectively (Supplementary Fig. 3). The optical fibers are arranged to ensure uniform illumination across the entire FOV (Supplementary Fig. 4). The entire system is enclosed in a distal cover with a front-facing sapphire window that allows direct contact with tissue surfaces for imaging. This cover can be readily removed for disinfection (Fig. 2a). The fully integrated system has a compact form factor with 14 mm diameter and 7 cm length, comparable to the size of a pen (Fig. 2b).

The phase mask design and image reconstruction algorithm were jointly modeled and optimized using an end-to-end deep learning framework (Fig. 2c, Supplementary Fig. 5). The first layer of the network implements a physics-informed model that simulates the image formation process of a conventional microscope with a learnable phase mask at the Fourier plane (see Methods). This simulation accounts for both fluorescence and reflectance imaging modes, and models defocus across 21 discrete depths spanning the 500 µm DOF. Following the optical simulation layer, digital reconstruction is performed by a modified U-Net architecture, which we refer to as PrecisionNet. This network incorporates several enhancements compared to basic U-Net architecture, including residual blocks, a pyramid pooling module (PPM), and pixel-shuffle convolution layers to improve feature representation and image quality (see Methods). The training dataset used for end-to-end optimization includes microscopy images of a wide range of features from both fluorescence and reflectance imaging modalities to optimize dual-modality imaging with PrecisionView (see Methods). The phase mask height profile and the reconstruction algorithm were simultaneously optimized during end-to-end training to minimize the loss.

Following end-to-end training, the optimized phase mask was fabricated using two-photon photolithography (Supplementary Fig. 6) and integrated into the PrecisionView system (see Methods). Once the system was fully assembled, a one-time calibration was performed by capturing PSFs at different depths and spatial locations across the FOV (Fig. 2d). These experimentally measured PSFs were then used to fine-tune PrecisionNet for improved reconstruction accuracy (see Methods). Compared to a conventional system with identical optics but without the trained phase mask, the captured PSFs of PrecisionView exhibited significantly improved contrast and consistency across the 500 µm DOF, both at the center and edges of the FOV. The modulated PSFs facilitate the network digital layer to perform image reconstruction and extend the FOV and DOF of the system (Supplementary Fig. 7).

### PrecisionView system characterization

We characterized the performance of PrecisionView using standardized test targets and fluorescent bead samples and found that the system achieves a lateral resolution of approximately 4 µm across a 500 µm DOF, with uniform performance maintained throughout the entire FOV. To ensure a fair comparison, we fabricated a reference system, referred to as Conventional in Figures 2, 3 and 4, using the same optical, illumination and housing components as PrecisionView, but without the phase mask or the reconstruction algorithm.

**Fig. 3.**
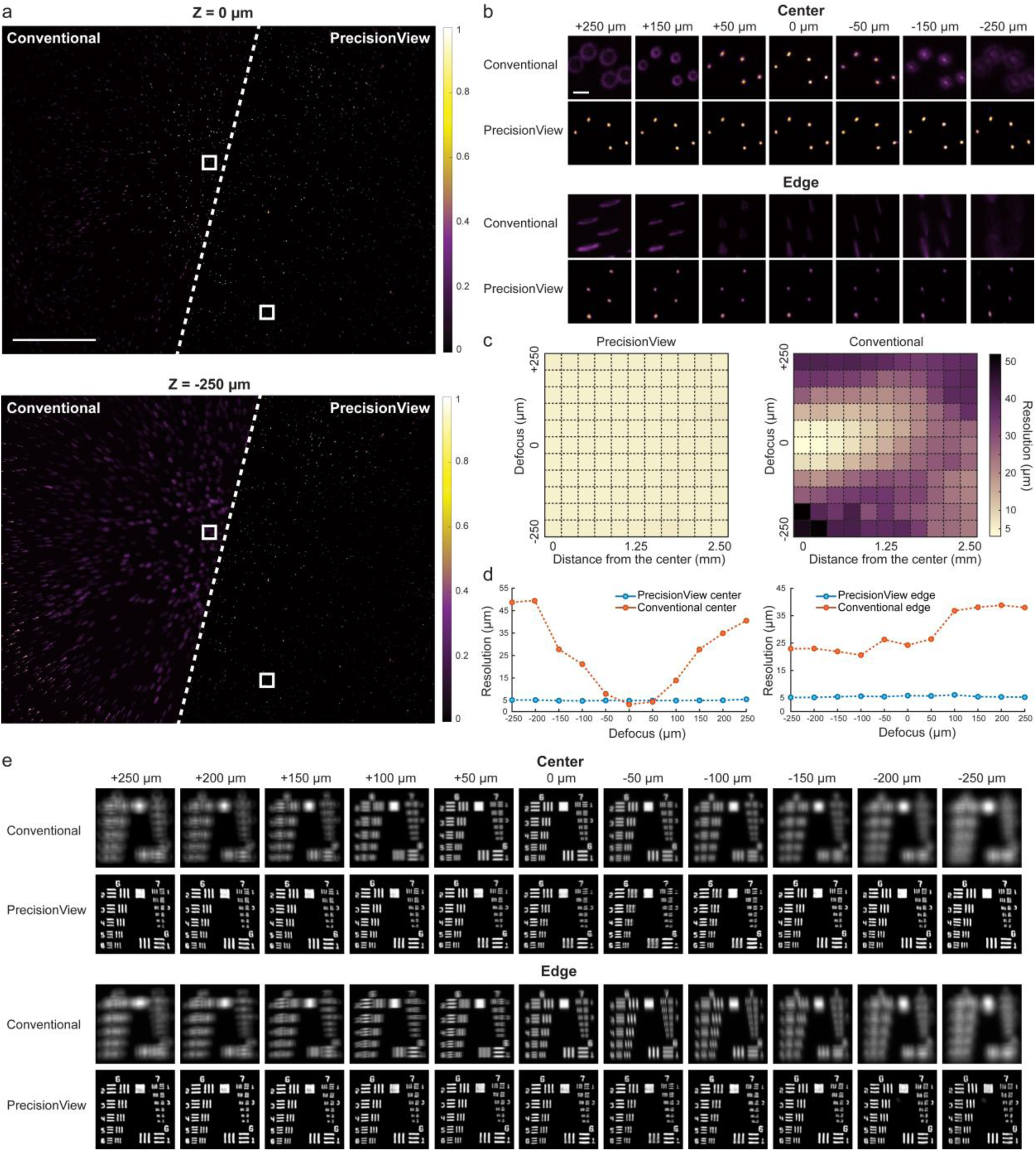
Characterization of the imaging performance of PrecisionView compared to a conventional system. **a** Experimentally captured images of 4 μm diameter fluorescent beads using PrecisionView and the conventional system at imaging depths of 0 μm and - 250 μm. Scale bar, 1 mm. **b** Zoom-ins of the boxed regions in panel a across a 500 μm depth range. PrecisionView maintains high-resolution visualization of beads consistently across this depth range, both at the center and edge of the FOV. Scale bar, 50 μm. **c** Experimentally measured lateral resolution across the entire DOF and FOV of PrecisionView and the conventional system. PrecisionView achieves a consistent lateral resolution of ∼ 4 μm, significantly outperforming the conventional system. **d** Comparison of the lateral resolution of PrecisionView and the conventional system across a 500 μm depth range, evaluated at the center (left plot) and edge (right plot) of FOV. **e** Experimentally captured images of a USAF test target using PrecisionView and the conventional system across a 500 μm depth range at the center and edge of the FOV. PrecisionView consistently resolves group 7, element 1 (3.91 μm line width).

**Fig. 4.**
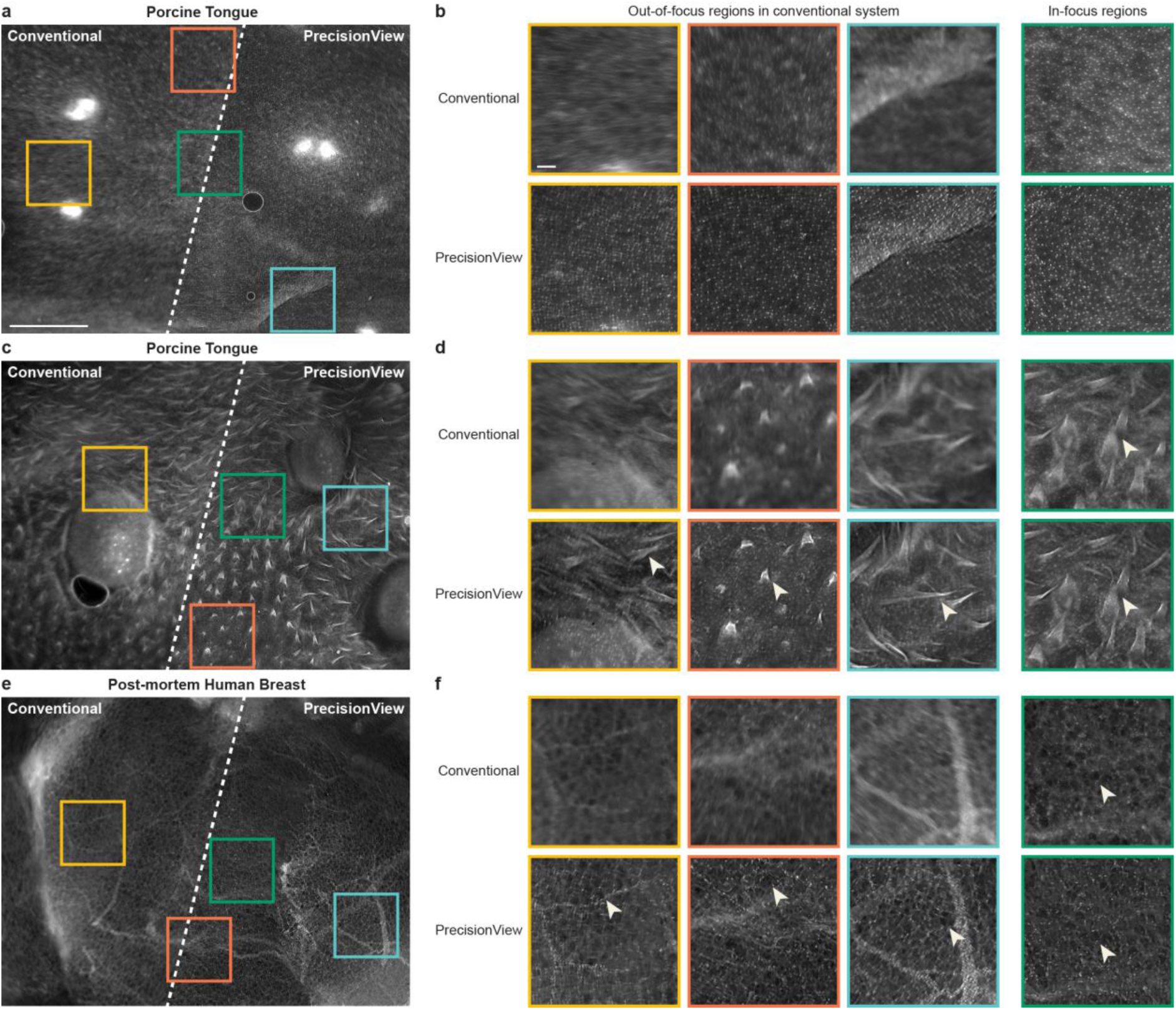
Representative ex vivo images of fresh porcine tongue and human breast specimens acquired with PrecisionView and the conventional system. **a** Representative images of a porcine tongue specimen. Scale bar, 1mm. **b** Zoom-ins of annotated ROIs in panel a. Scale bar, 100 μm. **c** Representative images of a different porcine tongue specimen showing distinct structural features. **d** Zoom-ins of annotated ROIs in panel c. Arrows indicate representative papillary features visualized within the selected ROIs. **e** Representative images of a post-mortem human breast specimen. **f** Zoom-ins of annotated ROIs in panel e. Arrows indicate representative adipocytes visualized within the selected ROIs. Results demonstrate that PrecisionView consistently resolves distinct cellular and epithelial morphology with high resolution across a large FOV in various tissue types, while the same features are blurred in various regions of FOV in images acquired with the conventional system due to geometric aberrations or limited DOF. In tissue regions that are in focus in the conventional system (green ROIs), the same cellular features are clearly resolved when imaged with PrecisionView.

We first imaged a glass slide with 4 µm diameter fluorescent beads spread across the slide using both PrecisionView and the conventional system (Fig. 3a). At the focal plane, both systems produced sharp, in-focus images at the center of the FOV. However, the conventional system showed noticeable image degradation toward the edges of the FOV, while PrecisionView maintained consistent image quality across the entire FOV. When imaging at defocused depths, only PrecisionView was able to recover clear images at both central and peripheral regions (Fig. 3a, b), highlighting its superior depth invariance and reduced sensitivity to optical aberrations.

We quantitatively evaluated the lateral resolution of PrecisionView and compared it to the conventional system across the full FOV and DOF. We imaged a glass slide with 1 µm diameter fluorescent beads spread across the slide, and the full width at half maximum (FWHM) of the bead intensity profiles was calculated at various spatial and depth positions for both systems (see Methods). PrecisionView maintained a consistent lateral resolution of approximately 4 µm throughout the 500 µm depth range and across the entire 20 mm² FOV, demonstrating its robust performance in both spatial and axial dimensions (Fig. 3c, d). In contrast, the conventional system exhibited high resolution only near the center of the FOV due to geometric aberrations and within a limited axial range of approximately 60 µm. Beyond this range, the lateral resolution of the conventional system degraded rapidly. These results highlight PrecisionView’s superior ability to maintain high-resolution imaging over approximately a 5-fold expanded FOV and 8-fold extended DOF, enabled by the optimized phase mask and computational reconstruction.

We further validated the performance of PrecisionView by imaging a negative 1951 USAF resolution target and comparing results to those from the conventional system (Fig. 3e). As expected, the conventional system was able to resolve the target elements at the focal plane; however, significant defocus blur appeared as the target was translated axially away from the focal plane, and image quality degraded toward the edges of the FOV. In contrast, PrecisionView consistently resolved Group 7, Element 1 (3.91 µm line width) across the entire 500 µm DOF, both at the center and edges of the FOV. These results further confirm the ability of PrecisionView to maintain high-resolution, in-focus imaging over an extended imaging volume.

### Ex vivo imaging of porcine tongue and human breast specimens

PrecisionView consistently resolves cellular features in freshly excised tissue specimens with high resolution across a large FOV. When imaging the same tissue regions, PrecisionView outperformed the conventional system without end-to-end optimization (Fig. 4).

For these experiments, fresh porcine tongue and post-mortem human breast tissues were cut with a scalpel and topically stained with 0.01% (w/v) proflavine solution in phosphate-buffered saline (PBS) using cotton-tipped applicators. The specimens were then placed on a glass window, and the system without distal cover was used to image from below to visualize the stained tissue surface through the glass window. For comparison, the same tissue regions were imaged without refocusing using the PrecisionView system and the conventional system configured with an identical optical layout, but without the phase mask. Fluorescence imaging was enabled using a blue LED light source for both systems.

Representative images from both tissue types are shown in Fig. 4, with three regions of interest (ROIs) selected per image for zoom-in visualization. We stained and imaged the squamous epithelium of the porcine tongue using both systems. Images of the squamous epithelium of porcine tissue acquired with PrecisionView clearly reveal cell nuclei with consistently high resolution across the entire FOV, while images acquired with the conventional system show blurred cellular features in portions of the FOV. (Fig. 4a, b). In regions of the porcine tongue with more complex features, PrecisionView effectively resolves multiple features including cell nuclei, keratinized epithelium, and papillae (Fig. 4c, d). In post-mortem human breast tissue, we imaged the cut surface of the sliced specimen. PrecisionView distinctly visualizes adipose tissue and cell nuclei, which appear poorly resolved in various regions of images acquired with the conventional system (Fig. 4e, f). In tissue regions that are in focus in the conventional system (green ROIs), PrecisionView accurately reveals the same cellular structures. These results demonstrate that PrecisionView enables in-focus, high-resolution imaging over a large FOV in fresh tissue specimens. Images acquired with PrecisionView consistently reveal epithelial and nuclear morphology across diverse tissue types, while images acquired with the conventional system show significantly blurred features in many regions within the FOV due to geometric aberrations or limited DOF.

### In vivo imaging of the oral cavity in healthy volunteers

We validated the in vivo imaging performance of PrecisionView in the oral cavity of healthy volunteers by visualizing cell nuclei and microvasculature in real time. The system was placed in gentle contact with the oral mucosa and scanned manually across large tissue areas while performing dual-modality imaging. With the large DOF, co-registered high-resolution images of cell nuclei and microvasculature across a large tissue area could be acquired without refocusing. As the system scanned the mucosal surface, the large FOV of each frame and the high video acquisition rate (∼ 15 FPS) provided substantial frame-to-frame overlap, enabling seamless stitching of dual-modality images to generate large-scale, co-registered maps (> 1.3 cm^2^) of the oral mucosa (Fig. 5). These maps clearly reveal distinct cellular and vascular features associated with anatomic variations with high spatial resolution.

**Fig. 5.**
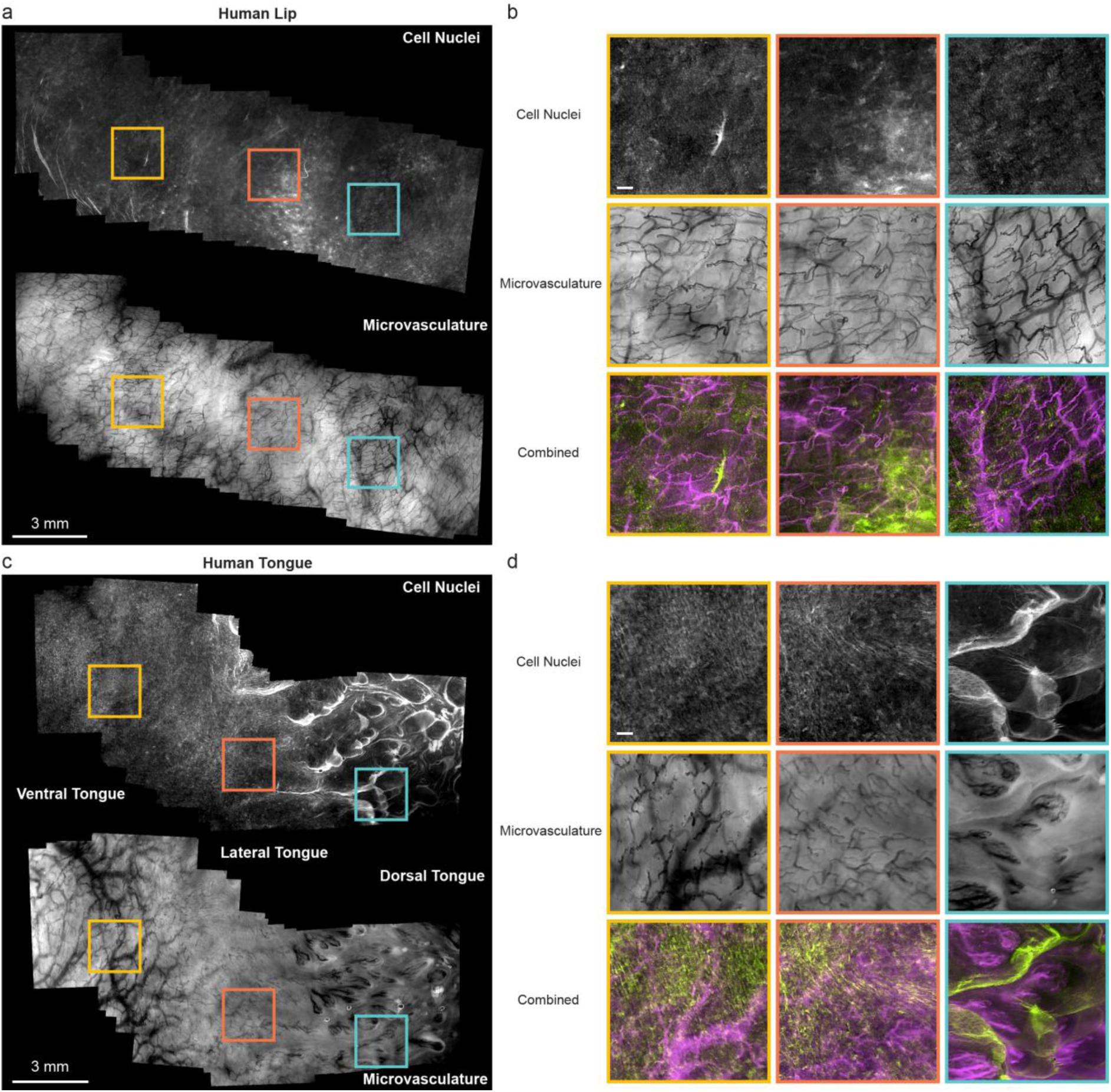
In vivo large-scale, high-resolution, co-registered mapping of cell nuclei and microvasculature in oral cavity of healthy volunteers using handheld PrecisionView imaging. **a** Stitched maps of cell nuclei and microvasculature of the lip mucosa of a healthy volunteer, covering a tissue area of ∼ 1.3 cm^2^. Video acquisition time, ∼ 1 min. Scale bar, 3 mm. **b** Zoom-ins of annotated ROIs in panel a, showing separate and combined high-resolution images of cell nuclei and microvasculature for each ROI. Scale bar, 200 μm. **c** Stitched maps of cell nuclei and microvasculature of the tongue of a healthy volunteer, covering a tissue area of ∼ 1.3 cm^2^. Video acquisition time: ∼ 1 min. Scale bar, 3 mm. **d** Zoom-ins of annotated ROIs in panel c, showing separate and combined high-resolution images of cell nuclei and microvasculature for each ROI. PrecisionView enables clear visualization of distinct tissue structures across the ventral, lateral and dorsal tongue. Scale bar, 200 μm.

Before imaging, tissue regions of interest in the oral cavity were topically stained with 0.01% (w/v) proflavine in PBS using cotton-tipped applicators. While imaging, we placed the PrecisionView system in gentle contact with the oral mucosa and manually scanned the stained tissue regions. The blue LED and green LED were alternated at 1 Hz to acquire fluorescence images of cell nuclei and reflectance images of microvasculature during scanning. Leveraging the system’s large FOV and extended DOF, co-registered high-resolution images were acquired without refocusing and stitched to generate co-registered maps of cell nuclei and microvasculature (Supplementary Video 2). Microvascular blood flow is clearly observed in several vessels as shown in the video. Representative stitched images of the lip mucosa covering a tissue area of ∼ 1.3 cm^2^ are shown in Fig. 5a. In the zoomed-in views of annotated ROIs (Fig. 5b), co-registered nuclear and microvascular structures are clearly visualized at high resolution. The extended DOF allows simultaneous acquisition of in-focus images of both cell nuclei and microvasculature. The large-scale maps reveal localized variations in the architecture of the microvasculature. The microvasculature map acquired from regions around the yellow and orange ROIs in the lip predominantly reveal capillary loops, while the microvasculature maps acquired from the blue ROI show a higher density of branching vessels.

Scanning across the ventral, lateral, and dorsal tongue epithelium with the PrecisionView system further demonstrated its ability to reveal distinct, localized tissue features at high resolution (Fig. 5c, d, Supplementary Video 3). On the ventral tongue (yellow ROI), both capillary loops and larger vessels are visualized with clearly resolved cell nuclei. The lateral tongue (orange ROI) shows similar features of cell nuclei and smaller vessels compared to the ventral tongue. While fluorescence imaging alone is not able to reliably distinguish between ventral and lateral tongue regions, the additional contrast from reflectance imaging significantly enhances the ability to differentiate these anatomic locations, highlighting the potential of dual-modality imaging. In images from the dorsal tongue (blue ROI), PrecisionView shows keratinized epithelium in fluorescence images and capillary loops embedded within the keratinized tissue in reflectance mode. The combination of co-registered fluorescence and reflectance images clearly reveal the microscopic structure of papillae in dorsal tongue.

### Ex vivo imaging of cervical specimens with precancerous lesions

To validate the capability of PrecisionView to detect precancerous lesions, we performed imaging on freshly resected human cervical specimens with squamous intraepithelial lesions (precancer). Following manual scanning of whole specimens and image stitching, dual-modality maps of the epithelium of the entire cervical specimen were generated for high-resolution assessment of epithelial and vascular structures (Fig. 6, Supplementary Video 4). Co-registered maps of the cell nuclei and microvasculature clearly delineated key anatomic structures, including columnar and squamous epithelium, as well as the squamocolumnar junction (SCJ). High-grade squamous intraepithelial lesions were clearly distinguishable from surrounding benign tissue, demonstrating the clinical potential of PrecisionView for noninvasive detection of neoplasia.

**Fig. 6.**
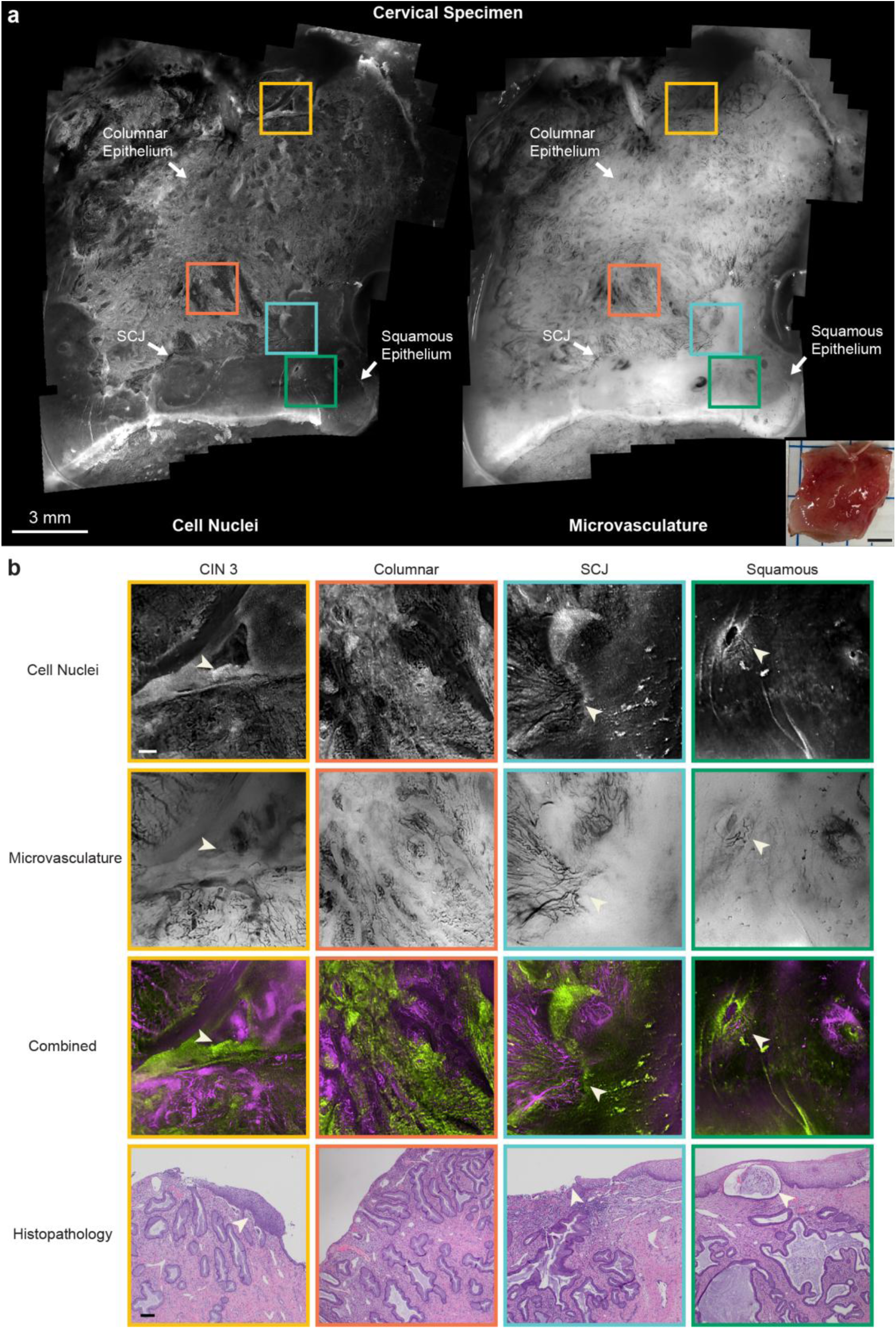
Ex vivo co-registered maps of cell nuclei and microvasculature from a freshly resected cervical specimen with precancerous lesions using PrecisionView. **a** Stitched dual-modality maps of cell nuclei and microvasculature from the epithelial surface of a freshly resected cervical specimen, covering a tissue area of ∼ 2.8 cm^2^. The co-registered maps delineate the columnar epithelium, the squamocolumnar junction (SCJ), and squamous epithelium. photograph of the imaged specimen is shown at the bottom right. Video acquisition time, ∼ 5 min. Scale bar of stitched maps, 3 mm. Scale bar of specimen photograph, 5 mm. **b** Zoom-ins of annotated ROIs in panel a, showing separate and combined high-resolution images of cell nuclei and microvasculature for each ROI. PrecisionView reveals a focal high-grade squamous intraepithelial lesion (CIN 3) in the background of columnar epithelium (indicated by arrows). The system also distinguishes columnar and squamous epithelium and accurately identifies the SCJ (indicated by arrows). An endocervical gland can also be observed within the green ROI located in the normal squamous region (indicated by arrows). Corresponding histopathology from same tissue regions confirmed the findings from PrecisionView. Scale bars, 200 μm.

In this pilot study, cervical specimens were obtained via loop electrosurgical excision procedure (LEEP) or cold knife cone (CKC) biopsy performed for the treatment of cervical dysplasia (precancer). PrecisionView imaging was carried out immediately after excision. Prior to imaging, the epithelial surface of ectocervical specimens was first cleaned with sterile swabs and saline to remove residual debris, then topically stained with 0.01% (w/v) proflavine solution in PBS. During imaging, the handheld PrecisionView system was placed in gentle contact with the stained tissue surface and scanned manually across the entire specimen. The blue LED and green LED were alternated at 1 Hz to enable dual-modality imaging during tissue scanning. Acquired images were stitched to produce high-resolution, co-registered maps of nuclear and microvascular features. Representative stitched images of a cervical specimen are shown in Fig. 6. Additional results are shown in Supplementary Fig. 8. Similar to the in vivo oral imaging, PrecisionView enabled simultaneous high-resolution visualization of both cell nuclei and microvasculature. The large-scale dual-modality maps and corresponding zoom-ins clearly delineate the columnar and squamous epithelium and highlight the SCJ. As shown in Fig. 6b, glandular structures and dense stromal vasculature are clearly visualized in the orange ROI within the columnar epithelium. The combination of nuclear and vascular features reveals the architecture of endocervical glands. In contrast, images of the normal squamous epithelium show uniform stratified cell nuclei and sparse capillary loops. An endocervical gland can also be observed within the green ROI located in the squamous region (indicated by arrows in Fig. 6b). The SCJ is distinctly visualized as the transition boundary between columnar and squamous epithelium (indicated by arrows in Fig. 6b). Imaging findings were confirmed by histopathological analysis of the same tissue regions. The extended DOF of PrecisionView enabled consistent, in-focus imaging across the entire tissue surface without the need for refocusing, despite significant variations in the microvasculature depth across different anatomic sites^20^. Notably, an island of suspicious squamous mucosa located within the columnar tissue region was highlighted in the dual-modality maps and corresponding zoom-ins (indicated by arrows in Fig. 6b). The suspicious squamous tissue showed dense nuclei, enhanced fluorescence, and increased vascular density compared to normal squamous tissue. Histopathological analysis confirmed the region as a high-grade squamous intraepithelial lesion (CIN 3). These results demonstrate the potential of PrecisionView for noninvasive, high-resolution imaging of large tissue epithelial surfaces, supporting its applications as a point-of-care diagnostic tool for detecting precancer and early cancer.

## Discussion

In this study, we present a compact, affordable ($3,000), AI-powered endomicroscope capable of high-resolution, dual-modality imaging of cell nuclei and microvasculature with a large FOV (> 20 mm^2^) and extended DOF (500 µm) for in vivo precancer and cancer detection. By integrating a deep learning-optimized phase mask and co-trained reconstruction algorithm, PrecisionView achieves a ∼5-fold improvement in FOV and ∼8-fold increase in DOF compared to conventional systems. This enables high-throughput, in-focus imaging of irregular tissue surfaces and simultaneous visualization of cell nuclei and microvasculature located at different depths. Through manual scanning with rapid video acquisition and image stitching, PrecisionView enables co-registered mapping of epithelial cell nuclei and microvasculature across large tissue surfaces with microscopic resolution. These results reveal distinct morphological features associated with anatomic structures and precancerous lesions, demonstrating the potential of PrecisionView as a powerful diagnostic tool for improving early neoplasia detection at the point-of-care.

Conventional IVM systems face fundamental trade-offs between resolution, FOV, DOF and system form factor. Achieving the high resolution required for pathology-level assessment and compact form factor necessary for clinical usability results in a limited FOV and small DOF. This restricts the ability to comprehensively evaluate large tissue regions of clinical suspicion, limiting the ability to provide important spatial context and determine which tissue regions might require biopsy and histologic assessment^11^. Other methods have been used to extend the DOF by enabling axial scanning using an electrically tunable lens^53–55^ or a deformable mirror^56^. However, these approaches increase system complexity and cost and are difficult to implement in handheld imaging systems due to their sensitivity to motion. In contrast, PrecisionView achieves significant improvement in both FOV and DOF through a simple optical layout and compact design, which are otherwise unachievable with conventional optics. The PrecisionView system can be readily assembled using off-the-shelf components and a customized phase mask at low cost. This hardware–software co-optimization strategy, powered by deep learning, provides new strategies for the development of next-generation IVM systems to overcome the intrinsic barriers that have hindered their broad clinical adoption. Extended DOF is achieved with each image capture, facilitating handheld clinical use. Unlike most computational microscopy techniques that rely on extensive post-processing, PrecisionView integrates rapid deep learning reconstruction directly into the video acquisition pipeline. This enables real-time image reconstruction and immediate visualization of in-focus microscopic features to facilitate point-of-care clinical evaluation. Although the camera can support video acquisition rates of up to 30 FPS, the effective acquisition frame rate is primarily limited by the long exposure time (∼65 ms) required for fluorescence imaging, corresponding to a frame rate of approximately 15 FPS in most experiments when performing dual-modality imaging. When operating only in reflectance mode for imaging microvasculature, PrecisionView can readily achieve faster video acquisition at 30 FPS. The real-time reconstruction speed is determined by the computation capability of the hardware. The PrecisionNet achieves real-time reconstruction at 22 FPS on a workstation with an RTX 4090 GPU, and at 7 FPS on a laptop with an RTX 3080 GPU. The customized GUI allows raw image acquisition at the maximum frame rate while displaying reconstructed frames at a lower rate.

Although image mosaicking has been used to expand the FOV of conventional IVM systems, the total stitched imaging area remains limited (< 10 mm^2^) due to the small FOV of each individual image acquisition and challenges in maintaining continuous scanning without abrupt positional shifts^21,34,35^. In contrast, with the ∼5-fold increase in FOV, PrecisionView significantly reduces sensitivity to abrupt shifts during manual scanning and enables acquisition of high-resolution images covering orders of magnitude larger tissue areas without missing regions. Moreover, the dual-modality capability of PrecisionView allows precise, co-registered detection of abnormal changes of both cell nuclei and microvasculature across large tissue surfaces to enhance clinical performance. To generate co-registered maps, we employed a stop-and-go acquisition pattern to scan the tissue (Supplementary Videos 1-4) which minimized the motion artifacts and ensured the acquisition of co-registered images of cell nuclei and microvasculature at each tissue site. Alternating between the two imaging modes at 1 Hz facilitates the real-time visualization and assessment of both nuclear and vascular features. With large-scale co-registered dual-modality maps, PrecisionView reveals detailed tissue architecture with high resolution, such as tongue papillae and endocervical glands. PrecisionView also accurately delineates various tissue anatomical subregions, which is critical for improving cancer detection. For example, since most squamous cell carcinomas of cervix arise in the transformation zone^57^, the ability of PrecisionView to clearly outline the SCJ enables more accurate assessment of high-risk regions and potential for improved identification of precancerous lesions. In addition, large-scale dual-modality maps facilitate the detection of focal abnormalities and distinct tissue structures that may otherwise be missed by conventional imaging systems with small FOVs and a single imaging modality.

With the ability for multiscale dual-modality evaluation of tissue, PrecisionView has the potential to serve as a powerful diagnostic tool for providing histopathological-level information directly at the point-of-care. PrecisionView has the potential to facilitate patient triage by determining which patient with a positive screening test needs a biopsy, provide highly precise guidance for biopsy location, and enable accurate surgical margin assessment at the point-of-care. It could help clinicians and surgeons to make immediate decisions such as whether additional resection is needed for positive margins, potentially reducing the need for on-site pathology or frozen section which is not available in many clinics and hospitals globally.

PrecisionView is designed to operate with minimal computational power. We used small-size images (368 × 480 pixels) for network training, allowing the model to be trained on standard workstations equipped with a GPU with 8 GB of memory. After one-time training, the trained network and customized user interface are deployed on a laptop, maintaining full-speed raw image acquisition and video-rate reconstruction. The frame rate of reconstruction is dependent on hardware specifications. PrecisionView has the potential to improve early detection for a variety of epithelial cancers, including cervical, oral, and anal cancer. The modular and versatile design of PrecisionView allows for broad imaging applications with minimal system modification. For example, the system can be adapted to use alternative contrast agents such as methylene blue, a dye with FDA approval for use in chromoendoscopy, to image cell nuclei with reflectance imaging^58^. The largest contributor to the system size of our current prototype is the off-the-shelf CMOS sensor, and future designs can be further miniaturized by reducing the size of the sensor-supporting circuitry using a customized printed circuit board (PCB). Our pilot study of ex vivo imaging in cervical specimens with precancerous lesions demonstrated PrecisionView’s ability to differentiate precancerous from benign tissue. Future in vivo clinical studies with larger sample sizes are warranted to rigorously assess the diagnostic performance of PrecisionView with statistical significance. With more clinical data, computer-aid diagnostic algorithms can be developed to automatically interpret PrecisionView images. This can further reduce the need for trained medical personnel, which is particularly valuable for improving cancer detection and early intervention in low-resource settings.

## Methods

### Optical design and device fabrication

The PrecisionView prototype was built utilizing a commercially available camera (XIMEA MU050MR-SY) with a monochrome Sony CMOS imaging sensor (IMX675 with 5 MP and 2.0 μm pixel size). The optical system of PrecisionView consists of two optical subassemblies, an emission filter (Chroma ET520/40m), and a phase mask, all mounted in 3D-printed housings (Formlabs Form 3+, Black Resin). The objective subassembly contains three achromatic lenses (#45-424, #49-658, #49-660, Edmund Optics), and the tube lens subassembly is identical to the objective lens and placed immediately after the emission filter and phase mask (Fig 2a, Supplementary Fig 2).

Illumination for the system was delivered via 14 plastic optical fibers (500 μm diameter, #02-532, Edmund Optics) arranged symmetrically around the imaging module and angled at 75 degrees to provide uniform illumination across the entire FOV (Supplementary Fig 4). The measured excitation power at the designed working distance was approximately 0.15 mW/mm^2^ for blue light and 0.05 mW/mm^2^ for green light, resulting in a maximum illumination intensity at the skin of less than the threshold limit value specified by National Conference of Governmental Industrial Hygienists (ACGIH). All fibers are bundled and coupled to an LED-based illumination module comprising a blue LED (Mouser LZ4-40B208-0000) with an excitation filter (#84-705 Shortpass Filter, Edmund Optics), a green LED (Mouser LZ4-40G108-0000_G2) with a bandpass filter (Chroma ET520/40m), and a dichroic mirror (Thorlabs DMLP490R) to enable switching between blue and green illumination (Supplementary Fig 3). An Arduino (UNO R3) was used to synchronize the illumination.

After assembly, the entire system was enclosed with a 3D-printed cap (Formlabs Form 3B, BioMed Black Resin), featuring a front-facing anti-reflection coated sapphire window (#20-633 Edmund Optics) for direct contact imaging. To maintain sterility and protect the device, the 3D-printed cap was disinfected following a standard high-level disinfection procedure using Cidex OPA (Advanced Sterilization Products) after each imaging session^59^.

### End-to-end training framework

A physics-informed deep-learning network was developed to simultaneously optimize the phase mask design and the image reconstruction pipeline. This end-to-end framework includes a learnable optical layer followed by a reconstruction module, referred to as “PrecisionNet”, which performs the final image recovery. The phase mask height map and reconstruction algorithm were simultaneously optimized during end-to-end training to minimize the loss.

### Learnable optical layer

In the end-to-end deep neural network, the learnable optical layer simulates the microscope’s optical imaging while incorporating passive phase modulation at the Fourier plane. Within this system design, both fluorescence and reflectance imaging in the green spectrum are modeled as incoherent imaging at a wavelength of 530 nm.

To accurately simulate the system, we employed a model based on Fourier optics theory^36,37^. Specifically, the phase mask was placed at the Fourier plane of the imaging system, where the point spread function (PSF) can be expressed as the squared magnitude of the Fourier transform of the pupil function.

The pupil function *P*(*x*_1_, *y*_1_, *z*) characterizes the spatial distribution of amplitude and phase modifications introduced to the wavefront at the Fourier plane:

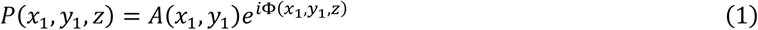

The amplitude *A* is defined by a circular aperture function, representing a disk-shaped aperture with values of unity inside the aperture and zero outside. The phase Φ in this system was modeled as the sum of two distinct components: a defocus component Φ^*DF*^, and learnable component representing the phase modulation introduced by the phase mask Φ^*M*^. The resulting phase Φ of the pupil function:

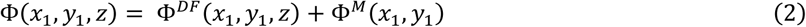

The defocus component Φ^*DF*^represents the defocus due to the mismatch between in-focus depth *z*_0_ and the actual imaging depth *z*. This can be modeled as:

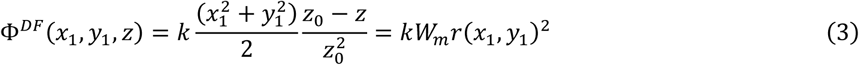

Where 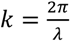 is the wavenumber and *λ* is the wavelength, 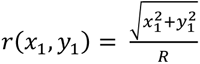 represents the normalized radial displacement in the Fourier plane, which is the radial coordinate normalized by the lens aperture radius *R*, and *W*_*m*_ denotes the maximum optical path-length error at the edge of the pupil caused by defocus.

The phase modulation introduced by the height variation on the phase mask Φ^*M*^ can be modeled as:

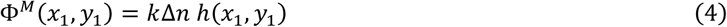

Where Δ*n* is the refractive index difference between air and the phase mask material, and ℎ represents the height map of the phase mask. The height map ℎ is a learnable parameter within the network and is iteratively updated during the training process.

The PSF in the imaging plane *PSF*(*x*_2_, *y*_2_, *z*) can be expressed as the squared magnitude of the Fourier transform ℊ{·} of the pupil function *P*(*x*_1_, *y*_1_, *z*):

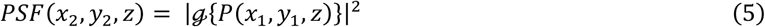

The image formation process can be described mathematically as the convolution of the object intensity distribution *I*_0_ with the system’s PSF:

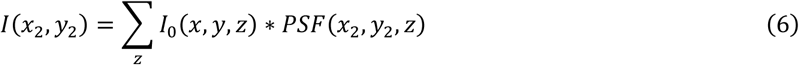

During the phase mask optimization step, we discretized the depth range into multiple layers, each blurred by the corresponding depth-specific PSF. We simulated image formation across a defocus range of −250 µm to +250 µm, discretizing this range into 21 distinct imaging depths. The discrete Fourier transform (DFT) was utilized to model the system’s PSF, computed using matrices of 25 × 25 pixels, with the discretization based on the sensor’s pixel size. To accelerate convergence during optimization while allowing adequate degrees of freedom, we further constrained the learnable height map by representing it in terms of a finite set of Zernike polynomial basis functions. Specifically, the height map ℎ was constrained as follows:

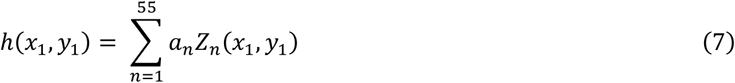

Where *Z*_*n*_(*x*_1_, *y*_1_) is the set of Zernike polynomials, and *a*_*n*_ is the coefficient vector. These coefficients were optimized during end-to-end training.

### PrecisionNet

Recovering an all-in-focus image from a blurry measurement without prior defocus information represents a blind deblurring problem, optimally addressed using deep neural networks. To solve this, we developed a deep neural network, we referred to as PrecisionNet, designed specifically for the image reconstruction task. PrecisionNet employs a modified U-Net architecture^60^ that leverages multi-scale feature extraction with skip connections between corresponding encoder and decoder blocks^61–63^. At each scale, PrecisionNet incorporates an additional residual block consisting of two convolutional layers with kernel size 3×3. Additionally, a pyramid pooling module (PPM)^64^ is integrated at the bottleneck layer to effectively capture global contextual information. The encoder utilizes pixel-shuffle convolution^65^ for downsampling, whereas the decoder employs bilinear interpolation for upsampling. The detailed architecture of PrecisionNet is illustrated in Supplementary Fig. 5. For memory-efficient training, PrecisionNet takes blurry input images sized 368×480 pixels and outputs the corresponding sharp, in-focus images. Once trained, PrecisionNet can reconstruct images directly at the raw capture resolution (1960×2592 pixels).

### Training dataset and implementation

To fully leverage the power of deep learning, a large and diverse set of training images that capture a wide range of imaging features is essential. However, collecting large-scale, paired datasets for lensless imaging systems remains a significant challenge, as supervised training typically relies on well-labeled ground truth data. To address this, we employed a forward model simulator to generate synthetic measurements based on various image sources, including open-source microscopy datasets^66–68^, histopathology images^37^, widefield fluorescence images of standard microscope slides acquired with a 4× (Nikon Fluor) objective^49^, proflavine-stained oral cancer resections imaged with a 10× (RMS10X) objective^37^, and widefield microscopy data containing vascular features^48^. Specifically, we selected 1,000 images from the open-source microscopy datasets, 600 histopathology images of healthy and cancerous human tissues (brain, lungs, mouth, colon, cervix, and breast) from The Cancer Genome Atlas (TCGA) FFPE slides, and 800 widefield microscopy images captured in-house for the training process. The training dataset includes microscopy images from both fluorescence and reflectance imaging modalities to optimize the dual-modality imaging of PrecisionView. From the combined dataset, we randomly selected 2,000 images for training, 200 for validation, and reserved another 200 for testing. During training, images were randomly cropped and augmented with rotations, flips, and brightness adjustments to improve model generalization.

Appropriate loss functions are critical for ensuring high-quality image reconstructions. The loss function used to train the end-to-end network was a weighted combination of pixel-wise L1 and L2 losses, computed between the reconstructed all-in-focus image stack and the corresponding ground truth images.

The network was trained using the Adam optimizer^69^, with a learning rate of 1e-9 for the optical layer and 1e-4 for the PrecisionNet. Training and testing were accelerated using a graphics processing unit (GPU). Specifically, model training was performed on a single GPU (Nvidia GeForce RTX 4090 GPU, 24 GB memory), requiring approximately 24 hours to complete 100 epochs with a batch size of 20. After training, the network achieved a reconstruction time of approximately 0.025 s per 8-bit frame at a resolution of 1960 × 2592 pixels on the same GPU. A custom user interface was developed to enable real-time visualization of both raw image captures and reconstructed output from the PrecisionView system. The interface supports real-time display of reconstructed video streams at a frame rate of up to 22 FPS on the training workstation, providing responsive feedback during imaging sessions. The user interface and trained reconstruction model were deployed on a laptop (Dell Mobile Precision with an Intel i9-11950H 8-core CPU and NVIDIA GeForce RTX 3080 GPU), enabling raw image acquisition at up to 30 FPS and real-time reconstruction at up to 7 FPS.

### Phase mask fabrication

The phase mask was fabricated using a 3D maskless two-photon photolithography system (Nanoscribe, Quantum X) in high-resolution dip-in liquid lithography mode (Supplementary Fig. 6). The mask was fabricated on a 700 μm thick fused silica substrate using the photoresist resin IP-S. The two-photon grayscale lithography enabled a smooth surface quality, while the adaptive voxel size could be tuned based on laser power variations, with a maximum power of 50 mW, a slicing distance of 1 μm, and a hatching distance of 0.2 μm. After exposure, the fabricated mask was immersed in SU-8 developer for 15 minutes, followed by a 2-minute soak in isopropyl alcohol. The substrate was laser cut to 12 mm diameter to fit the design of the housing and fitted with an opaque mask to create an aperture containing only the phase mask.

### Device calibration and network fine-tuning

For each assembled system, a one-time calibration procedure was performed to record the experimental PSFs of the system prior to imaging, accounting for differences between simulated and experimental PSFs caused by alignment and fabrication tolerances. The point source used for calibration was a 3 μm pinhole array with 500 µm spacing that was illuminated by a green LED (Thorlabs, M530L4) placed behind an 80-degree holographic diffuser. The pinhole array was first focused on the focal plane and scanned axially from −250 µm to +250 µm at a step size of 50 µm to form focal stacks, which were further used to generate training pairs. In total, 80 PSFs were captured per depth, and 11 depths were calibrated. All calibration images were averaged through ten captured images to improve the signal-to-noise ratio.

When the phase mask was incorporated into the system, we observed a significant reduction in spherical aberration. However, the PSFs continued to vary across the FOV, which degraded reconstruction performance when PrecisionNet was fine-tuned using only a single, centrally calibrated PSF. To address this issue, we utilized a comprehensive set of experimentally calibrated PSFs captured across different depths and spatial locations for network fine-tuning. During retraining, these experimentally measured PSFs were used in a forward model to generate simulated ground truth–capture image pairs. PrecisionNet was then trained on this dataset, allowing the network to learn and correct for PSF variations across the entire FOV. This approach enabled robust handling of spatially varying blur, significantly improving the consistency and quality of the reconstructed images.

### Fluorescent bead imaging and resolution characterization

To evaluate the imaging performance of PrecisionView, we prepared glass slides coated with 4 µm diameter (Molecular Probes FluoSpheres 4.0 μm, yellow-green fluorescent, 2% solids) and 1 µm diamter fluorescent beads (Fluoresbrite YG Microspheres 1.0 µm, Cat # 17154-10) and performed imaging with PrecisionView and the conventional system for comparison. Beads were imaged at different axial positions ranging from −250 µm to +250 µm from the focal plane with a step size of 50 µm. To quantitatively evaluate resolution, we calculated the FWHM of the intensity profiles of 1 µm beads at various lateral and axial positions for both systems. At each axial depth, 250 µm × 250 µm square ROIs were selected at different spatial locations within the FOV. The FWHM of intensity profiles were calculated along both horizontal and vertical directions for all visible beads within the ROI and averaged to quantify the local lateral spatial resolution. The same process was repeated at all depths for both PrecisionView and the conventional system.

### Animal specimen preparation and imaging

Freshly resected ex vivo porcine tongue specimens were obtained from a local abattoir. A small portion of tongue epithelium surface was excised with a scalpel and topically stained with 0.01% (w/v) proflavine solution in PBS using a cotton-tipped applicator. The stained tongue epithelium was first imaged using the PrecisionView system. For comparison, the same tissue regions were subsequently imaged using a conventional microscope configured with an identical optical layout, but without the phase mask.

### Post-mortem human specimen preparation and imaging

Post-mortem human breast tissue was acquired from Accio Biobank Online. A small portion of the tissue was cut with a scalpel and the tissue cut surface was topically stained with 0.01% (w/v) proflavine solution in PBS using a cotton-tipped applicator. The stained tissue surface was first imaged using the PrecisionView system. For comparison, the same tissue regions were subsequently imaged using a conventional microscope configured with an identical optical layout, but without the phase mask.

### In vivo human oral imaging

The in vivo human study was conducted at Rice University; study participants were adult volunteers aged 18 years or older and without underlying health conditions. The study protocol was approved by the Institutional Review Board (IRB) at Rice University, and written informed consent was obtained from each participant prior to imaging. Before each imaging session, the device underwent a standard high-level disinfection procedure^59^. Tissue regions of interest in the oral cavity were stained topically with a 0.01% (w/v) proflavine solution in phosphate-buffered saline, applied using cotton-tipped applicators. Imaging was performed immediately following proflavine application. During imaging, the PrecisionView system was gently placed in contact with various stained tissue sites within the oral cavity. Reconstructed microscopic images showing epithelial cell nuclei and subsurface microvasculature were displayed in real time via the graphical user interface (GUI), while raw image frames were saved for post-session analysis. Each imaging session lasted approximately 1 to 2 minutes, allowing examination of multiple tissue sites of interest. The acquired raw frames were subsequently processed using PrecisionNet and stitched into composite views using the Image Composite Editor (Microsoft). A total of six healthy volunteers were enrolled in this study. Consistent imaging quality was achieved across all healthy volunteers enrolled in the study.

### Surgical sample imaging

The study was conducted at The University of Texas MD Anderson Cancer Center (MD Anderson), with protocol approval from the IRBs of both MD Anderson and Rice University. Eligible participants included patients aged 18 years or older who were scheduled to undergo LEEP or CKC biopsy for the treatment of cervical dysplasia and had a confirmed negative pregnancy test. Written informed consent was obtained from all participants prior to enrollment. Before each imaging session, the device underwent a standard cleaning and high-level disinfection^59^. Immediately following excision, the cervical specimens were imaged using the PrecisionView system. The tissue surface was first cleaned with sterile swabs and saline to remove any residual debris, then stained topically with a 0.01% (w/v) proflavine solution in phosphate-buffered saline, applied using cotton-tipped applicators. During imaging, the PrecisionView system was held by hand and placed in gentle contact with the stained tissue surface, then scanned across the entire specimen. Reconstructed microscopic images revealing epithelial cell nuclei and subsurface microvasculature were displayed in real time via the graphical user interface (GUI), while raw image frames were saved for further analysis. Each imaging session lasted approximately 5 to 8 minutes, enabling comprehensive examination of entire excised cervical specimen. After PrecisionView imaging, the cervical specimens were sent to the pathologist within the expected time for histopathological diagnosis per standard of care. The raw image data captured by PrecisionView were subsequently processed using PrecisionNet and stitched into composite views using the Image Composite Editor (Microsoft). Nine patients were enrolled in this study, with consistent imaging quality achieved across all subjects. Data from two patients were shown in Fig. 6, Supplementary Fig. 8 and Supplementary Video 4. Imaging findings were correlated with final pathology results.

## Data availability

The main data supporting the results of this study are available within the paper and its Supplementary information. The raw and analyzed datasets generated during the study are available for research purposes from the corresponding author upon request. Source data is provided with the paper.

## Code availability

Custom codes used in this study are available on a GitHub repository at (https://github.com/JiminWu/PrecisionView).

## Acknowledgements

This work was supported in part by the National Institute of Dental and Craniofacial Research of the National Institutes of Health under Award Number R01DE029590 for R.R.R.-K. and A.M.G., Award Number R01DE032051 for R.R.R.-K., A.V., and A.M.G., and NSF grant IIS-1730574 and IIS-1652633 for A.V. Additional support was provided from the United States National Cancer Institute through the MD Anderson Cancer Center Support Grant P30CA016672 for K.M.S., P.R., A.M.G., and M.P.S., and U54CA277843 for K.M.S., R.R.R.-K., A.V., and M.P.S.. The content is solely the responsibility of the authors and does not necessarily represent the official views of the National Institutes of Health. This research is partially sponsored by the Defense Advanced Research Projects Agency (DARPA) through Cooperative Agreement D20AC00002 (for J.T.R. and A.V.) awarded by the U.S. Department of the Interior (DOI), Interior Business Center. The content of the information does not necessarily reflect the position or the policy of the Government, and no official endorsement should be inferred. The authors thank all the healthy volunteers and patients who volunteered to participate in the study. The authors also thank Andrew Kim and Cindy Melendez for coordinating patient recruitment and enrollment at the University of Texas MD Anderson Cancer Center, and Dr. Barrett Lawson from the Department of Pathology at the University of Texas MD Anderson Cancer Center for facilitating the clinical imaging workflow.

## Contributions

H.H. and J.W. contributed equally and either has the right to list themselves first in bibliographic documents. H.H. and J.W. designed, fabricated, and characterized the PrecisionView prototype, and performed the imaging experiments and data analysis. V.B. designed PrevisionNet architecture and provided guidance for the network training. J.L. and T.S.T. developed the phase mask fabrication process and fabricated the phase mask. A.S. developed the user interface for PrecisionView. H.H., K.G., A.M.G., M.P.S., and K.M.S. performed imaging of surgical specimens. A.M.G., P.R., M.P.S., and K.M.S. contributed clinical expertise and provided surgical specimens. J.C., R.A.S., J.T.R., A.V., and R.R.R.-K. provided guidance and assistance with all aspects of the work. All authors contributed to the writing of the manuscript.

